# Phenological, Physiological and yield markers as efficient tools to identify drought tolerant rice genotypes in Eastern India

**DOI:** 10.1101/2020.05.29.122929

**Authors:** Soumya Kumar Sahoo, Goutam Kumar Dash, Arti Guhey, Mirza Jaynul Baig, Madhusmita Barik, Selukash Parida, Padmini Swain

## Abstract

Rice production is severely threatened by drought stress in Eastern India. To develop drought tolerant varieties, selection of donors for breeding programme is crucial. Twenty one selected rice genotypes including both tolerant and sensitive to drought were grown under well-watered and drought stress conditions in dry seasons of two successive years of 2017 and 2018. Leaf water potential, relative water content displayed significant difference among the genotypes during vegetative screening. At reproductive stage drought screening, days to 50% flowering was delayed in all genotypes except N22 and Anjali (showed early flowering) however grain yield and other yield related traits decreased significantly compared to well watered condition. Correlation analysis of phenological and yield related traits with grain yield revealed that tiller numbers and panicle numbers are highly correlated with grain yield both under well-watered and water stress conditions and contributes maximum towards grain yield. The dendrogram grouped Mahamaya, Sahabhagidhan, Poornima, IBD 1, Hazaridhan, Samleshwari and Danteshwari into one cluster which performed better under water stress conditions and had grain yield more than 1.69 tha^−1^. Sahabhagidhan, Poornima, Vandana, and N22 displayed tolerance to drought both under vegetative and reproductive conditions which could be a good selection for the breeders to develop drought tolerant rice cultivars for eastern region of India.

## INTRODUCTION

Although rice grows better in sufficient water available conditions, yet it has better adaptability to highly diverse ecological condition. India is the centre of rice diversity (Ray *et al*., 2015) and the crop is cultivated in about 43 million hectares of land with 110 million tons of milled rice production was recorded as per 2016-17 statistical reports. About 40% of rice producing area of India is rainfed, out of which 70% is present in eastern India in the states of Odisha, Chhattisgarh, West Bengal, Bihar, Jharkhand and Eastern Uttar Pradesh. The total rainfed area of this region comprises 77% of lowland and 23% of upland out of which 52% of lowland and entire area of upland are affected by drought (Pathak *et al* 2018).). In the year 2017, India had approximately 43 m ha rice growing area (Keerlery 2020), nearly 60% of which were in Eastern India (IRRI, 1997) and only rainfed rice growing area of Eastern India accounts for 12.9 mha (Lal et al. 2017). These areas characterized with often failure of rain or a long spell between two rains, hence drought stress can appear at any period of crop growing stage that may be at seedling, vegetative, and reproductive stage or it can be intermittent drought depending upon the rainfall pattern and distribution. Although drought at reproductive stage is more detrimental, still drought at vegetative stage also a determining factor for reproductive growth in rice.

Water requirement of rice crop depends on field conditions, cropping seasons and growth stages (MARDI, 2009). Since, rice plants require water throughout their growth period, there are certain critical growth stages when drought stress can dramatically reduce the grain yield (Bajji *et al*., 2001; Zhou *et al*., 2007). During vegetative stage, drought stress decrease the leaf and tiller formation that ultimately reduce yield by affecting panicle development (Swain *et al*. 2017; Singh *et al*. 2017). However, when drought stress occurs during reproductive growth phase, it remarkably reduces rice grain yield due to abortion of ovule and formation of partially filled grains (Pantuwan *et al*. 2002a).

Drought stress tolerance can be potentially measured through physiological and phenological parameters. Plant metabolism primarily dependent on water status and the best way to determine plant water status is through measurement of relative water content (RWC) and leaf water potential (LWP). Maintenance of proper plant water status The relative water content (RWC) is crucial physiological parameter that refers to the degree of cell or tissue hydration which is responsible for maintaining growth activities that results in better grain yield in rice (Silva *et al*. 2007). Measurement of RWC indicates the stress intensity and act as a screening tool for plant water status (Hassanzadeh *et al*. 2009). Gutierrez *et al*. (2010) and Manickavelu *et al*. (2006) investigated on RWC, which is highly correlated with morphological and yield traits viz., plant height, days to flowering, panicle length and harvest index besides grain yield. Several reports showed that higher decrease in LWP was observed in drought susceptible genotypes than the tolerant ones (Silva *et al*., 2010; Silvestre *et al*., 2017). The lower leaf water potentials leads to reduced turgor, stomatal conductance and photosynthesis, and thus eventually reduce grain yield (Akbarian *et al*., 2011; Amini *et al*., 2014). Standard evaluation system (SES) score (drought score and drought recovery score) is expressed as an elective way to deal with plant drought tolerance (Fen *et al*. 2015). Drought score taken at seedling stage is commonly not a good yield determining index (Mitchell et al., 1998) but at later growth stages it is considered as an essential parameters for drought screening (IRRI, 2014). Drought score and drought recovery score measurement is a convenient way to determine the degree of tolerance to oxidative damage in plants and show lack of hydration of the plant tissue related with its RWC (Cabuslay *et al*., 1999; Cabuslay et al 2002). Zhu (2002) and Mishra (2005) documented that under drought stress shoot growth and plant height reduction is a very common morph-physiological adaptation to water stress and is crucial for survival practice. Water deficiency, reduces plant height and tillers number (Zhang *et al*., 2009). The number of tillers is also linked with leaf rolling score, drought score and proline accumulation. So, lower tiller production during water stress may be a determinant of drought tolerance (Fen *et al*., 2015).

Drought decrease rice biomass accumulation at vegetative stage (Zhang *et al*., 2018), that results in grain yield reduction due to lowered filled grain number at reproductive stage (Fukai *et al*., 1999). In rice, under drought low grain yield is mainly due to the reduced number of filled spikelets per panicle without a substantial change in the number of spikelets per panicle (EKANAYAKE *et al*., 1989; Wei *et al*., 2014). Liu *et al*. (2006) reported that flowering stage is more vulnerable to water limited condition than any other developmental stage. Water stress at the booting (Pantuwan *et al*., 2002) and flowering stages interfere in floret initiation, leading to reduction in number of panicles per plant, grain filling percentage, spikelet sterility and decreased 1000 grain weight (Nour *et al*., 1995 and Fabre *et al*., 2005) which lead to poor grain yield in rice (Acuna *et al*., 2008). In the present study, twenty one popular genotypes are evaluated both under vegetative and reproductive stage drought to identify suitable donors for breeding programme.

## MATERIALS AND METHODS

The experiment was conducted taking twenty one rice genotypes (including tolerant check Sahabhagidhan and susceptible check IR 20) to study their response towards drought stress both at vegetative and reproductive stage during dry season of 2017 and 2018 at the experimental field of ICAR-National Rice research Institute (ICAR_NRRI), Cuttack, Odisha (85°55’48’E-85°56”48”E and 20°26”35’N-20°27”20’N with the general elevation of 24m above the MSL). Out of twenty one genotypes, seven genotypes were native to Chhattisgarh and were collected from Gene bank of Indira Gandhi Krishi Vishwavidyalaya, Raipur and rest fourteen genotypes were collected from Gene bank of ICAR-NRRI.

### Plant growth and treatments

The genotypes were grown in field both under well-watered (WW) and drought stress (DS) conditions in randomised block design with three replications. Two to three seeds were sown per hill at a depth of 2cm with a spacing of 20 cm between rows and 15 cm in between hills. In both the growing seasons; well-watered plot was maintained with nearly 5cm standing water from 30 days of germination to till maturity. To impose water stress, two separate experiments were conducted: vegetative stage screening (VS) and Reproductive stage screening (RS) for two consecutive years 2017 and 2018.

Vegetative stage drought stress was imposed by withholding irrigation for a period of 30 days, when seedlings were 30 days old. After 30 days of stress period (60 DAG), drought stress was released by irrigating the experimental field. Relative water content (Schonfeld et al. 1988), and leaf water potential Turner (1988) was recorded at the end of the stress. Drought score and drought recovery score was recorded according to Standard Evaluation System for rice (IRRI, 2002).

To impose stress at reproductive stage, irrigation was withdrawn 10 days before flowering and surface irrigation was provided till when soil moisture content dropped below 17% and soil moisture tension dropped below −65kPa. To syncronize flowering dates, the genotypes were divided into four groups and staggered planting was adopted. Observations like days to 50% flowering, plant height, tiller number grain yield and yield attributes such as panicle number, total dry matter, harvest index, fertility percentage and 1000 test grain weight were recorded after maturity (Yoshida *et al*. 1981).

The soil of both the fields was sandy clay loam, moderately acidic pH (4.5–5.5) with medium organic carbon content (0.5–0.75%). The recommended fertilizers doses of N: P_2_O_5_: K_2_O @ 80: 60: 60 kg ha^−1^ in the form of urea, DAP and MOP, respectively was applied and plant protection measures were used as and when required during the crop growth period.

To assess the intensity of drought stress, soil moisture content (SMC) was measured using time domain refractometer (TDR) from a soil depth of 15cm and 30cm at an interval of 5 days after withdrawal of irrigation. Soil moisture potential (SMP) was observed daily from a soil depth of 15cm and 30cm using tensiometer and water table depth was measured installing piezometers in the experimental plots.

### Statistical analysis

The obtained phenological, physiological and yield related data were calculated and scatter plots were constructed by using Microsoft excel. Statistical comparison between variance was carried out by ANOVA (Analysis of variance) among the genotypes and the water treatments using CROPSTAT ver 7.2. Correlation among the different traits were determined by using SPSS package ver23. The dendrogram was constructed using XLSTAT evolution 2020.1.1.

## RESULTS AND DISCUSSION

The performance of the twenty one rice genotypes were evaluated under well-watered (WW) and drought stress (DS) conditions during dry seasons of 2017 and 2018. The results of the phenological, physiological and yield attributes were discussed on the basis of pooled data, obtained from results of 2017 and 2018.

### Soil moisture status as affected by drought stress

During drought stress period at the vegetative stage growth period (30-60 DAG), the SMC and SMP were decreased and WTD was dropped with the increase in the duration of stress period in both the dry seasons. During dry season 2017, at 36-38 DAG, the crop received a 55mm of rainfall and during dry season 2018, at 58 DAG the crop received 12mm of rainfall. (**Fig 1 & 2**). At 15 cm soil depth, the SMC decreased to 13.60% and 14.35% whereas SMP decreased to −51.50 kPa and −56.27 kPa during dry season of 2017 and 2018 respectively. At 30 cm of soil depth the SMC decreased to 15.60% and 17.75%, and SMP reached up to −47.83 kPa and - 54.55 kPa during dry seasons 2017 and 2018 respectively (**Fig 3 & 4**). WTD dropped below 102 cm and 97 cm in 2017 and 2018 respectively. From the SMC, SMP and WTD data, it is evident that the crop had experienced moderate to severe drought stress during the vegetative stage stress period.

**Fig 1.**
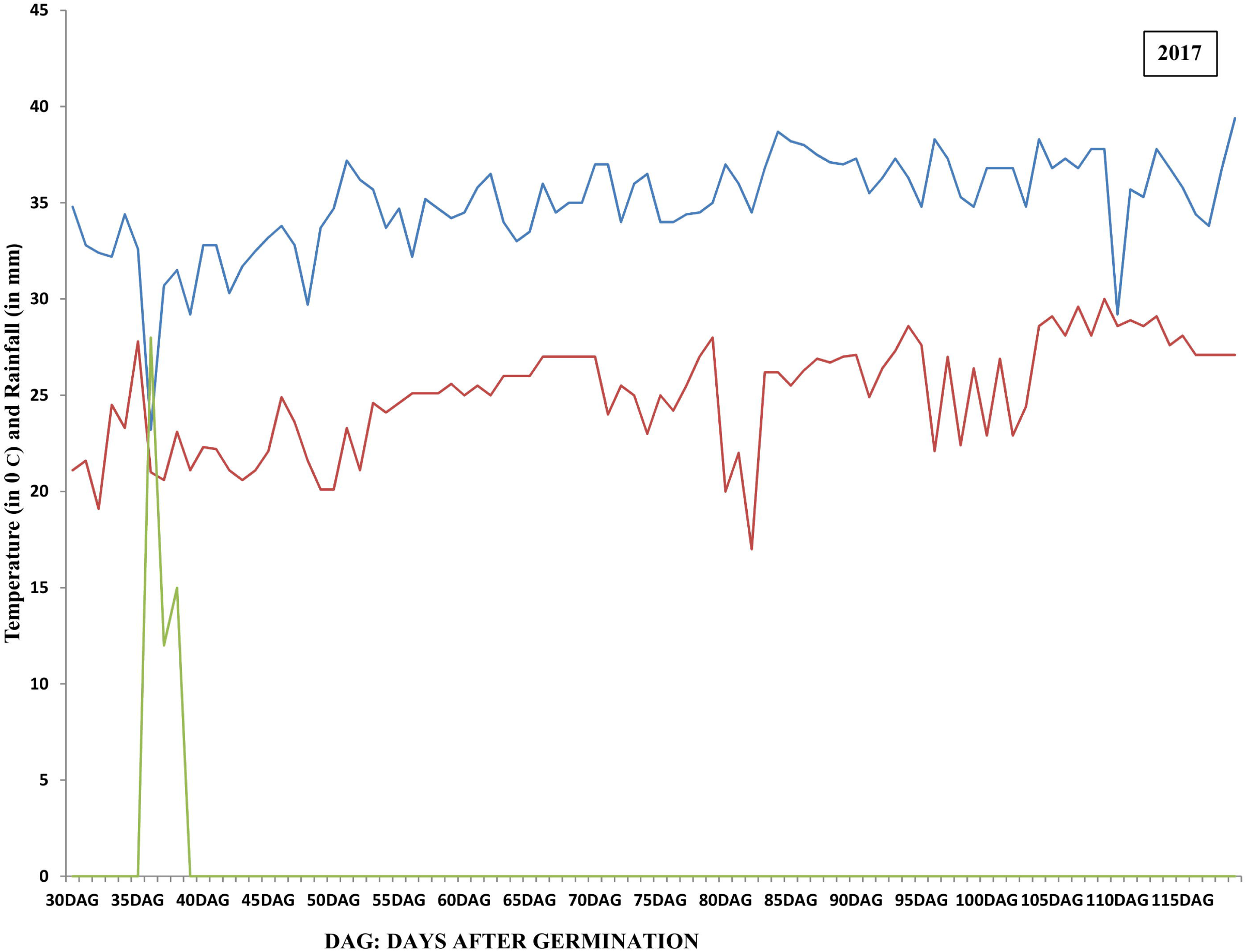
Meteorological data depicting maximum, minimum temperature (°C) and rainfall (mm) pattern during crop growth period of dry seasons 2017. DAG- days after Germination.

**Fig 2.**
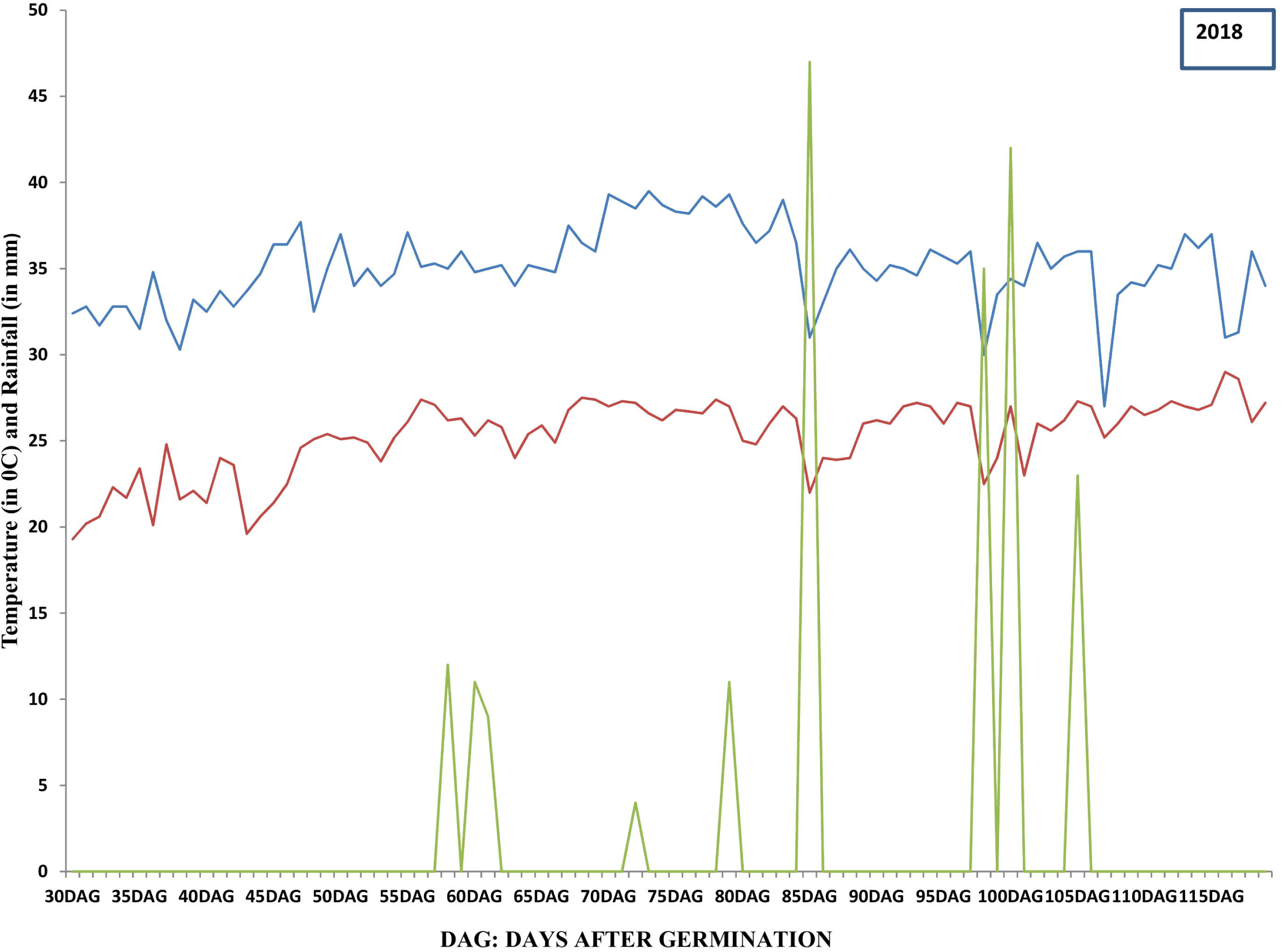
Meteorological data depicting maximum, minimum temperature and rainfall pattern during crop growth period of dry seasons 2018. DAG- days after Germination.

**Fig 3.**
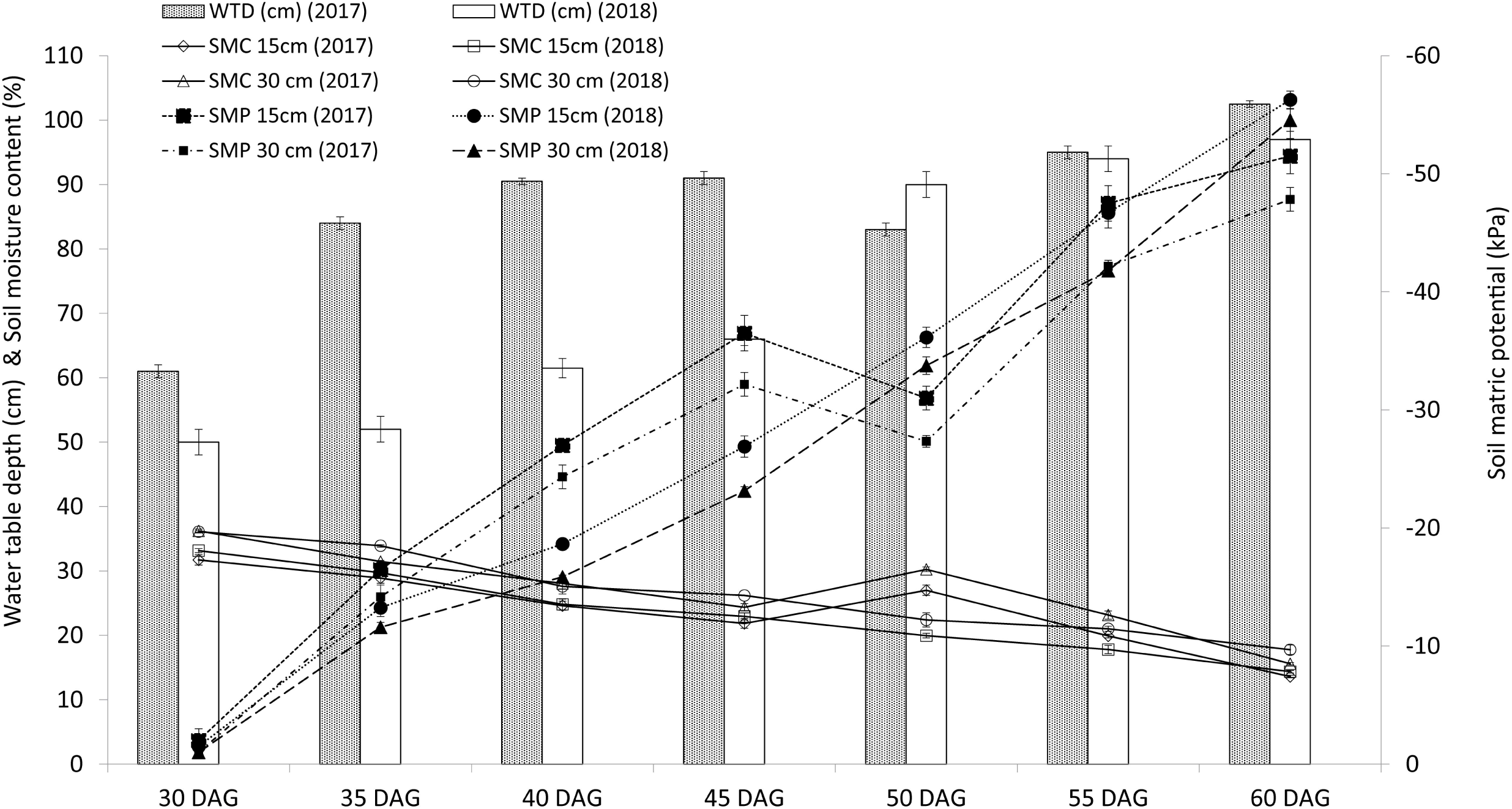
Soil moisture content (SMC %), soil matric potential (SMP -kPa) and water table depth (cm) during the vegetative stage drought stress period in dry season 2017 and 2018.

**Fig 4.**
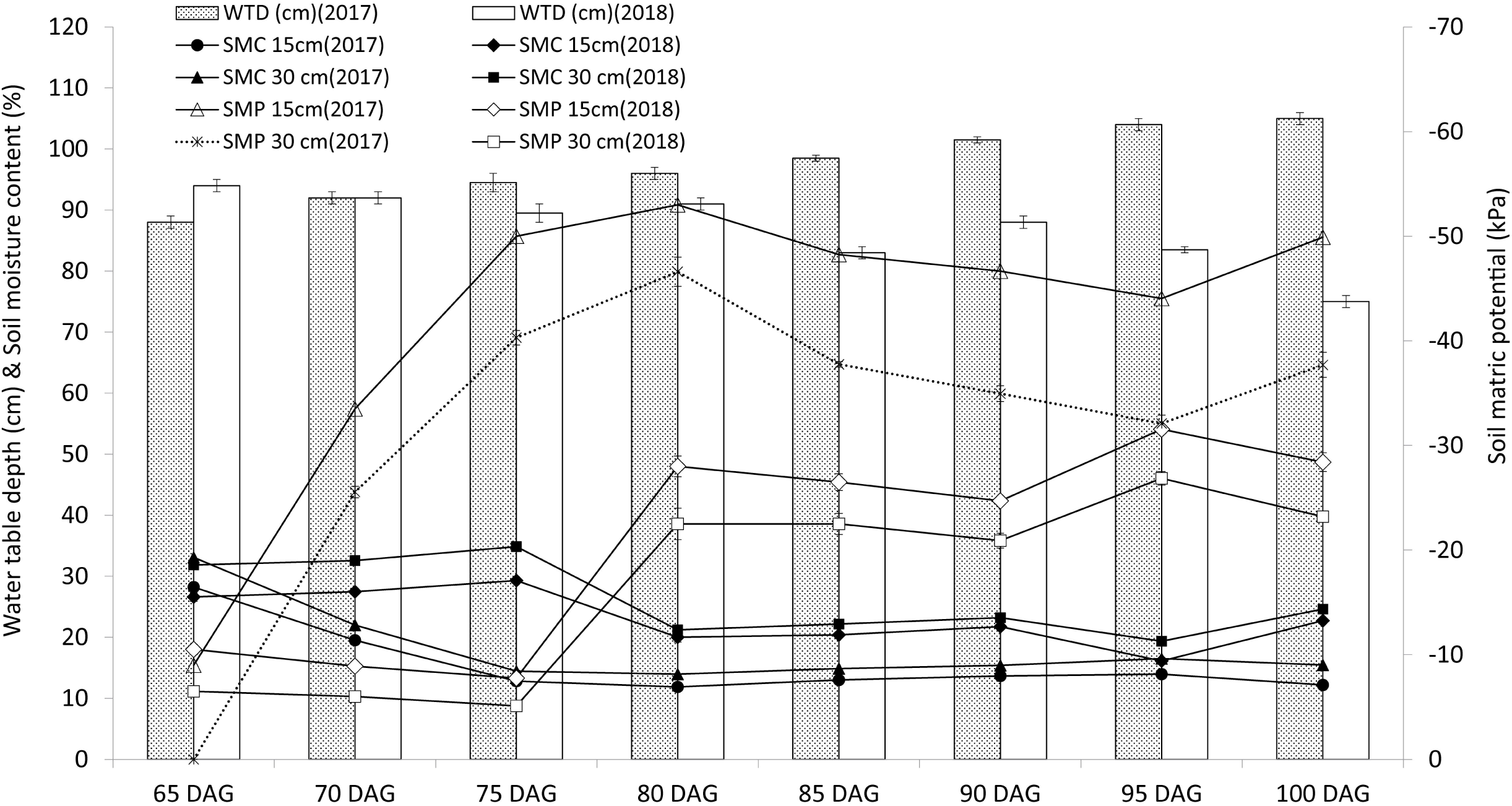
Soil moisture content (SMC %), soil matric potential (SMP -kPa) and water table depth (cm) during the reproductive stage drought stress period in dry season 2017 and 2018. DAG: days after germination.

**Table1.**
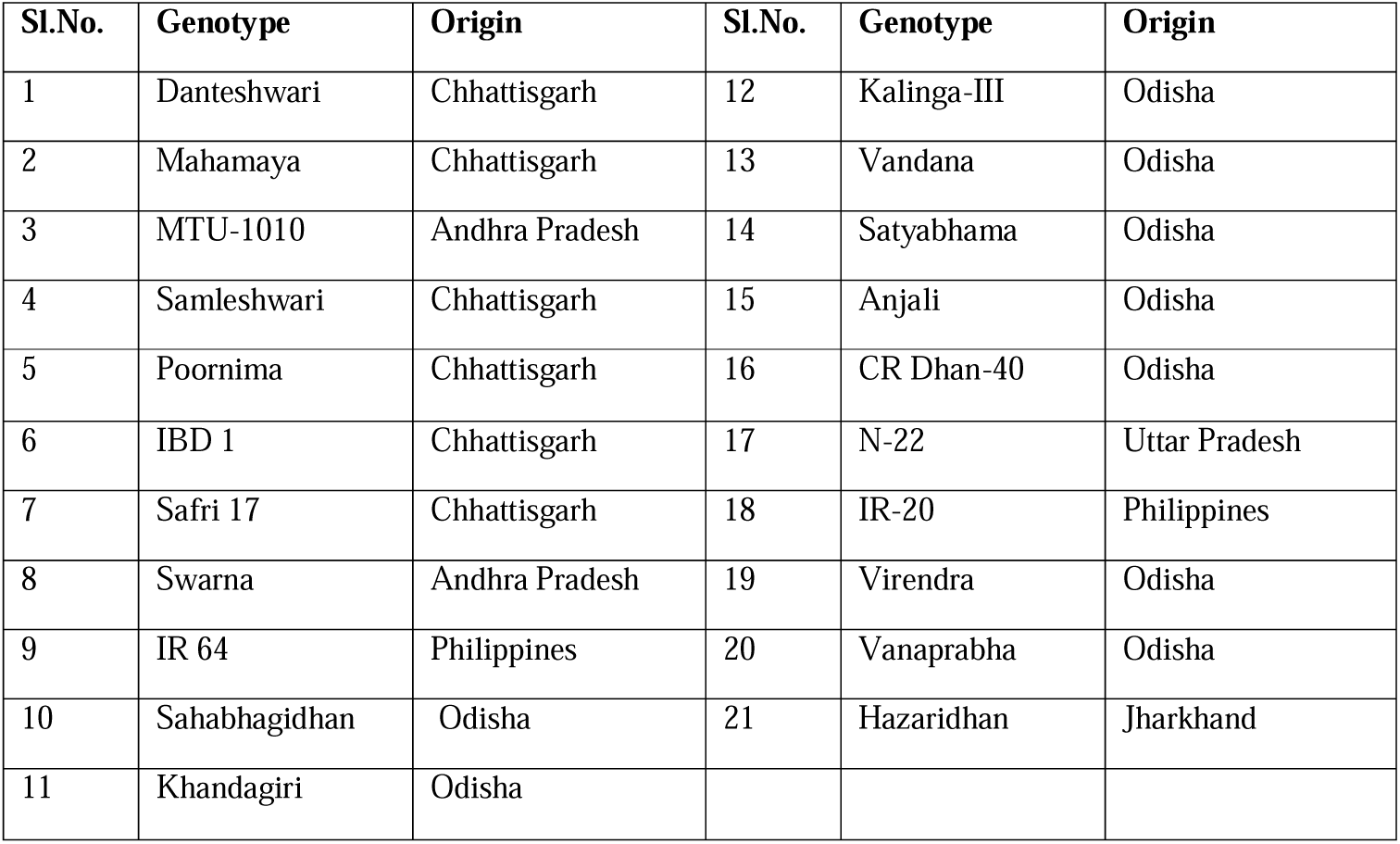
Information of the rice genotypes used in the experiment

### Phenological and Physiological traits as affected by water stress

In this study, the drought response of 21 rice genotypes for two years were evaluated based on SES score (**Table 2)**.

**Table 2.** Standard Evaluation System (SES) score for drought in dry seasons of 2017 and 2018.

| DRY SEASON-2017 |  |  |  | DRY SEASON-2018 |  |  |  |
| --- | --- | --- | --- | --- | --- | --- | --- |
| Sl. no | Genotypes | Drought Score | Recovery Score | Sl. no | Genotypes | Drought Score | Recovery Score |
| 1 | Danteshwari | 3 | 3 | 1 | Danteshwari | 3 | 3 |
| 2 | Mahamaya | 1 | 3 | 2 | Mahamaya | 1 | 3 |
| 3 | MTU1010 | 5 | 5 | 3 | MTU1010 | 5 | 5 |
| 4 | Samleshwari | 1 | 3 | 4 | Samleshwari | 1 | 3 |
| 5 | Poornima | 1 | 1 | 5 | Poornima | 1 | 1 |
| 6 | IBD-1 | 1 | 3 | 6 | IBD-1 | 1 | 3 |
| 7 | Safri 17 | 1 | 3 | 7 | Safri 17 | 1 | 3 |
| 8 | Swarna | 3 | 3 | 8 | Swarna | 3 | 3 |
| 9 | IR64 | 5 | 5 | 9 | IR64 | 5 | 5 |
| 10 | Sahabhagidhan | 1 | 1 | 10 | Sahabhagidhan | 1 | 1 |
| 11 | Khandagiri | 3 | 3 | 11 | Khandagiri | 3 | 3 |
| 12 | Kalinga-iii | 3 | 3 | 12 | Kalinga-iii | 3 | 3 |
| 13 | Vandana | 3 | 1 | 13 | Vandana | 1 | 1 |
| 14 | Satyabhama | 3 | 3 | 14 | Satyabhama | 3 | 3 |
| 15 | Anjali | 3 | 3 | 15 | Anjali | 3 | 3 |
| 16 | CR Dhan40 | 3 | 3 | 16 | CR Dhan40 | 3 | 3 |
| 17 | N22 | 1 | 1 | 17 | N22 | 1 | 1 |
| 18 | IR20 | 7 | 7 | 18 | IR20 | 7 | 7 |
| 19 | Virendra | 3 | 5 | 19 | Virendra | 3 | 5 |
| 20 | Vanaprabha | 3 | 5 | 20 | Vanaprabha | 3 | 5 |
| 21 | Hazaridhan | 3 | 5 | 21 | Hazaridhan | 3 | 3 |

In both the years, 7 genotypes (Mahamaya, Samleshwari, Poornima, IBD-1, Safri 17, Sahabhagidhan and N22) had SES drought score ‘1’, 11 genotypes (Vandana, Danteshwari, Swarna, Khandagiri, Kalinga III, Satyabhama, Anjali, CR Dhan 40, Virendra, Vanaprabha, Hazaridhan) had ‘3’ score, two genotypes (MTU 1010 and IR 64) had ‘5’ score and IR 20 had ‘7’ score in both the seasons, however, no genotypes had score ‘0’ and ‘9’ (**Table 2**). After 24 hrs of stress release, 4 genotypes (Sahabhagidhan, Poornima, Vandana, and N22) recovered with score ‘1’, 11 genotypes (Danteshwari, Mahamaya, Samleshwari, IBD-1, Safri 17, Swarna, Khandagiri, Kalinga III, Satyabhama, Anjali, CR Dhan 40) recovered with score ‘3’, 4 genotypes (IR 64, Virendra, Vanaprabha, Hazaridhan) recovered with score ‘5’ and IR20 did not recovered in both the seasons (**Table 2**).

Significant variation in LWP and RWC was observed between the treatments and among the genotypes (**Fig 5 and 6**). Mahamaya, Samaleswari, Poornima, IBD-1, Safri 17, Sahabhagidhan and N22 had higher LWP (>-3.50 MPa) with more than 70% leaf RWC under drought condition whereas MTU1010, IR 64 and IR 20 had lowest LWP (< - 4.5 MPa) with less than 60% RWC.

**Fig 5.**
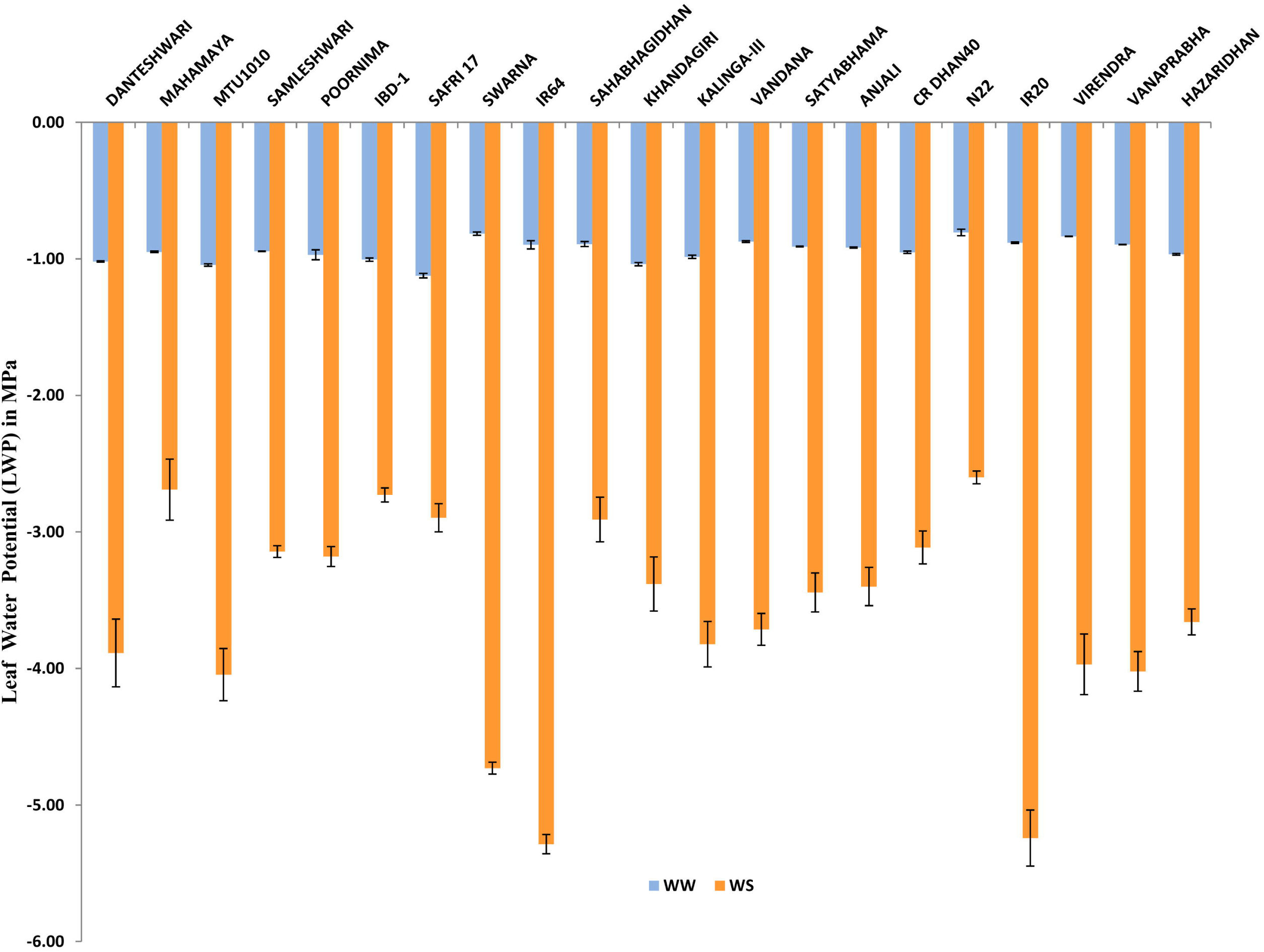
Pooled data of Leaf Water Potential (MPa) under well-watered and drought stress conditions of 21 genotypes

**Fig 6.**
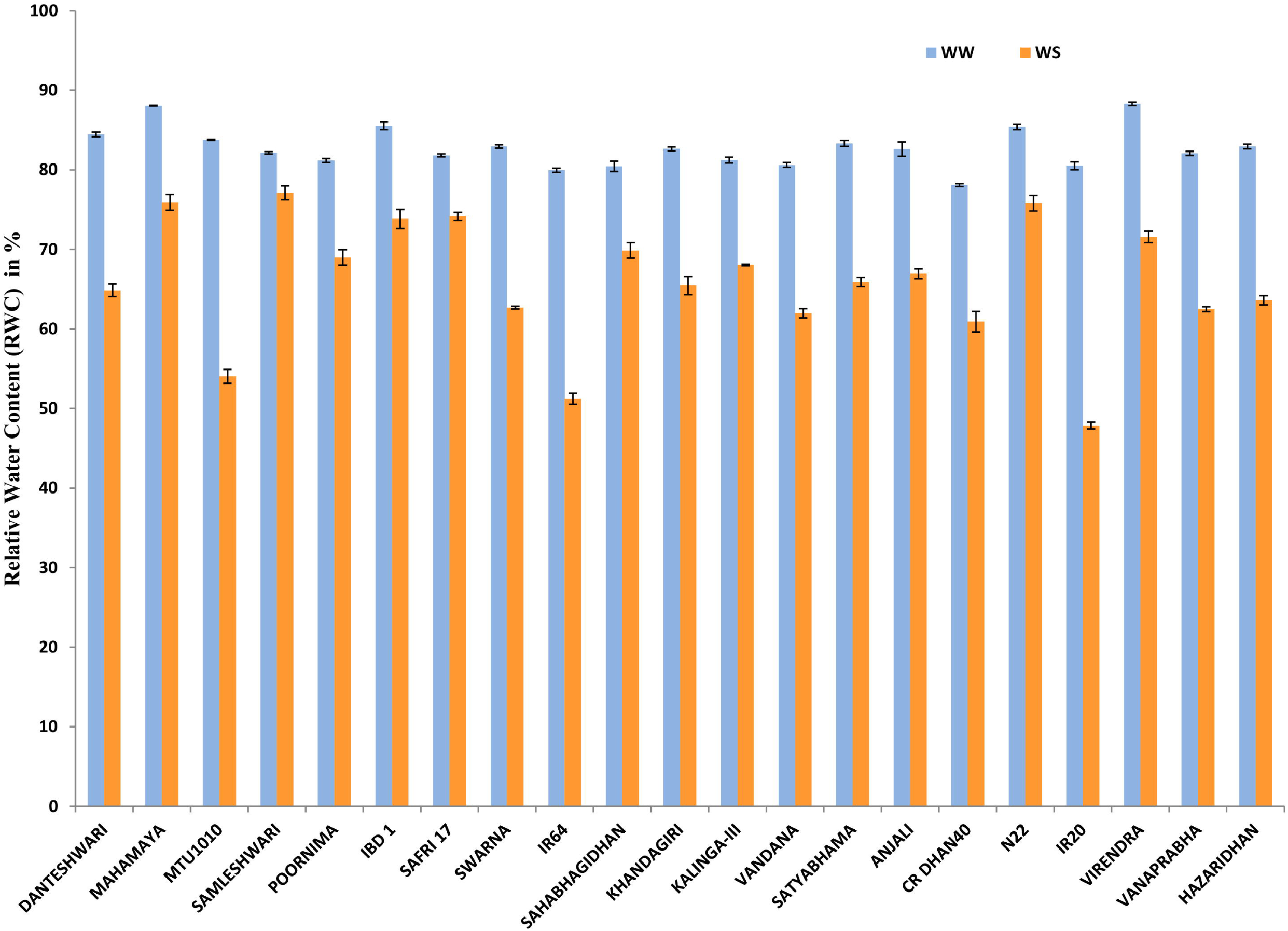
Pooled data of leaf relative water content (RWC %) under well-watered and drought stress conditions of 21 genotypes

In DS condition, maximum DFF was recorded in susceptible check IR-20 (112.33 days) followed by Swarna (104 days) and IR64 (96.00days), whereas N22 (59.17 days) took minimum days to attain flowering followed by Anjali (63.5 days) and Kalinga III (64.50 days) **(Fig 7)**. All genotypes except N22 and Anjali delayed in their DFF in DS as compared to WW condition however maximum delayed was observed in IR64 and IR-20 with 9.33 days (10.77% and 9.06%) followed by MTU1010 with 8.17 days (9.78%) and Swarna with 8.17days (8.52%). Minimum delayed in DFF was observed in Danteshwari and Vandana with 1.33days (1.87% and 2.00%.). As mentioned earlier Anjali and N22 flowered early by 7.17days (10.14%) and 5days (7.79%) respectively under DS condition.

**Fig 7.**
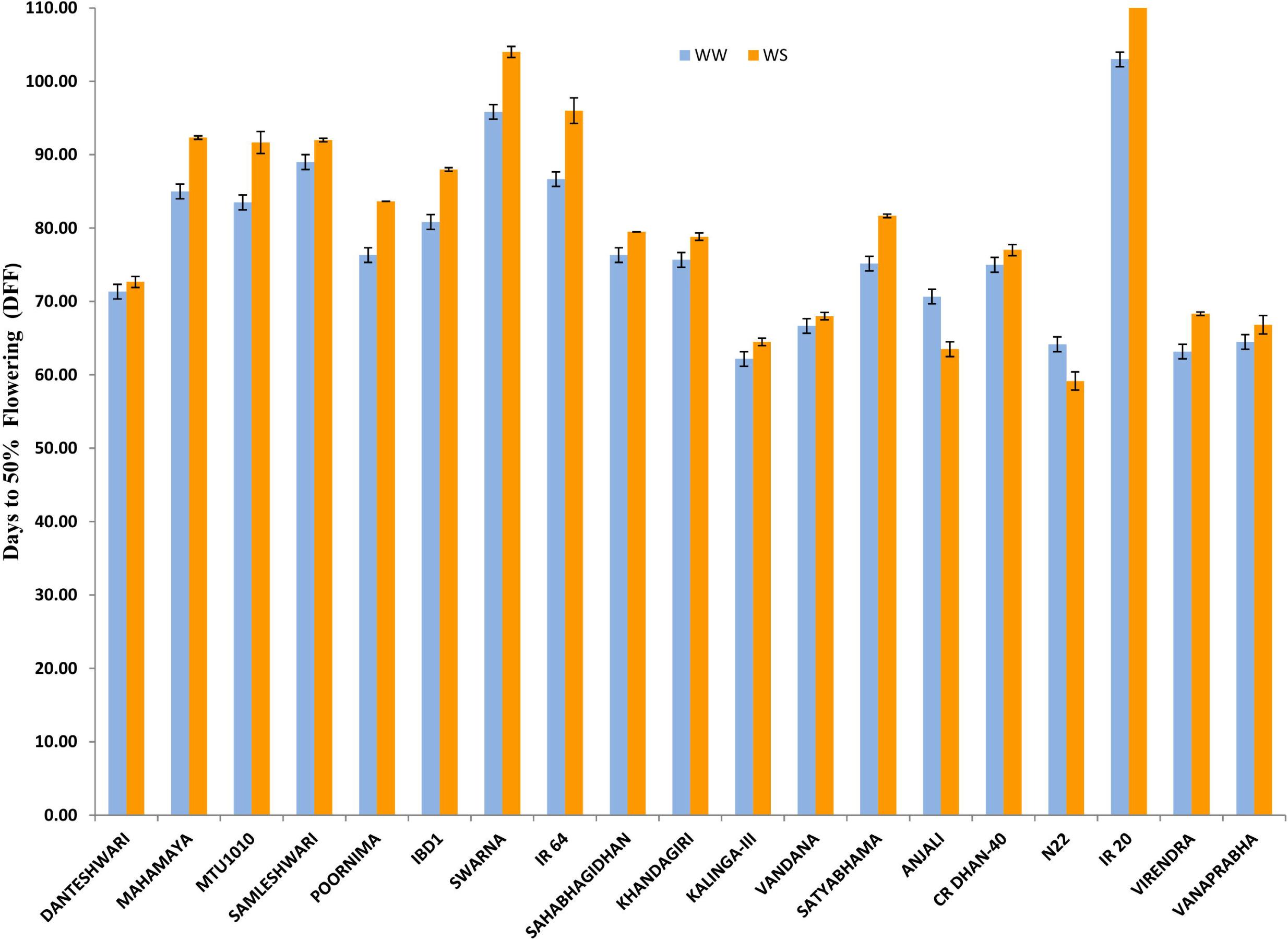
Pooled data of days to 50% flowering (DFF) data under well-watered and drought stress conditions of 20 genotypes

### Effect of water stress treatments on grain yield and yields attributes

Results of ANOVA (**Table 3**) indicated significant variation for the measured nine traits (PH, TN, PN, TDM, GY, RYR, HI, GF%, and GW) in dry seasons of both the year 2017 and 2018 among the genotypes and between the water treatments ranging from 3.39% for GW to 62.19% for GY. The pooled data of two dry seasons showed that grain yield suffered a mean yield loss of 62.19% (2.43 tha^−1^) and associated reductions in PH by 11.87% (12.07 cm), TN by 25.72% (77.64 no. per m^2^), PN by 32.68% (123.15 no. per m^2^), TDM by 54.80% (6.33 tha^−1^), HI by 19.15% (0.06), GF% by 43.33% (35.30%) and GW by 3.39% (0.76g) under DS as compared to WW condition.

**Table-3.**
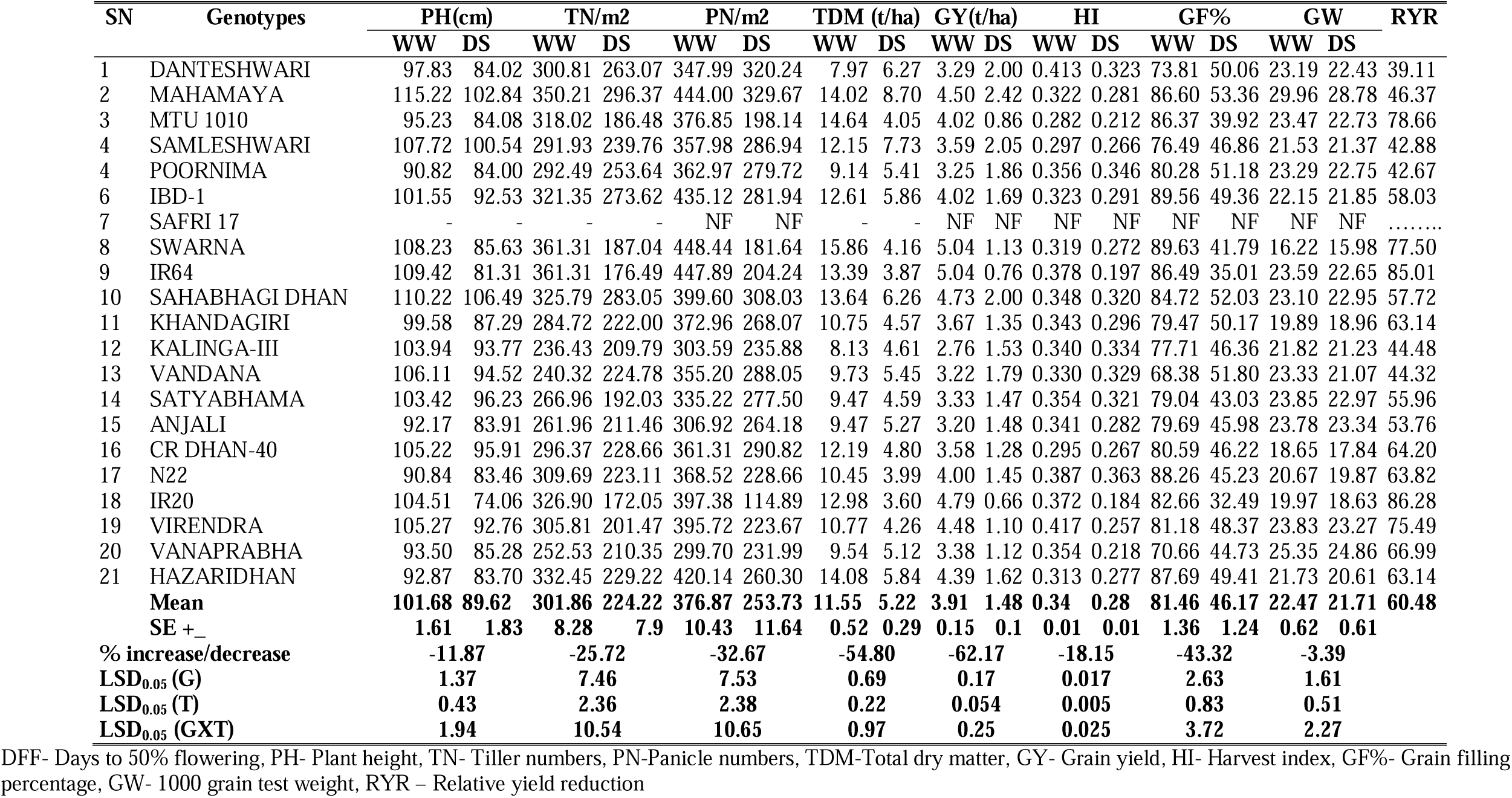
Grain yield and yield attributes of 21 rice genotypes under well-watered (WW) and drought stress (DS) of the dry season from pooled data of 2017 and 18.

### The performance of tested genotypes in response to drought stress

The results related to grain yield and yield attributes of rice genotypes under well-watered and drought stress conditions have been presented in **Table 3**. Under DS condition, higher grain yield was recorded in Mahamaya (2.42 t.ha^−1^), Samleshswari (2.05 t.ha^−1^), Sahabhagidhan and Danteswari (2.00 t.ha^−1^) while lowest grain yield was recorded in susceptible check IR20 (0.66 t.ha^−1^) followed by IR64 (0.76 t.ha^−1^) and MTU 1010 (0.86 t.ha^−1^). Minimum RYR was exhibited by Danteswari (39.11%), Poornima (42.67%) and Samleshwari (42.88%) while maximum RYR was observed in susceptible check IR20 (86.28%), IR64 (85.01%) and MTU1010 (78.66%) under DS condition over WW condition. The reduction in grain yield under DS condition resulted due to consistent and significant reduction in PH, TN, PN, TDM, HI, GF%, and GW in comparison to WW condition. The minimum reduction in PH was exhibited by tolerant check Sahabhagidhan (3.38%) followed by Poornima and Samleshwari while, maximum reduction was recorded in susceptible check IR-20 by 30.46 cm (29.14%) under DS as compared to WW condition. Similarly, the percentage reduction in TN was varied from 6.47% (Vandana) to 51.15% (IR64) under DS as compared to WW condition. Mahamaya, Danteshwari and Sahabhagidhan had PN >300 nos. m^−2^ whereas, susceptible check IR20 (114.89 nos. m^−2^) attained lowest value in DS condition with maximum percentage reduction (71.09%) compared to WW condition. Danteshwari followed by Anjali and Satyabhama showed least reduction (7.97-17.22 %) in PNm^−2^ in DS as compared to WW condition. Under DS condition Mahamaya, Samleshwari and Danteshwari had highest TDM (8.70 - 6.27 tha^−1^) while lowest was recorded by susceptible check IR20 (3.60 t.ha^−1^). Comparatively less reduction in TDM was observed in Danteshwari, Samleshwari and Mahamaya with 21.37%, 36.39% and 37.92% and higher reduction was recorded in Swarna, MTU1010 and IR64 with 73.79, 72.35% 72.29% respectively in DS condition as compared to WW condition. In DS condition, maximum HI was recorded in N22, Poornima and Kalinga III (>0.334) whereas, minimum HI was observed in IR20, IR64 and MTU 1010 (<0.212). Maximum reduction in HI was observed in IR20 (50.56%) while minimum was observed in Vandana (0.55%) in DS condition as compared with WW condition. Mahamaya, Sahabahagidhan and Vandana had higher GF% (>51.80) under DS condition while, susceptible check IR20 attained lowest GF% (32.49%). Comparatively lower percentage reduction in GF% was noticed in Vandana, Danteshwari and Poornima (24.25 - 36.25%) and higher percentage reduction was noticed in susceptible check IR20 (60.70%) in DS as compared to WW condition. In case of 1000 grain weight (GW), the maximum reduction was observed in Vandana (9.69%), IR20 (6.69%) and Hazaridhan (5.19%) while minimum reduction was recorded in Sahabhagidhan (0.68%), Samleshwari (0.76%) and IBD-1 (1.38%) in DS over WW condition. Minimum yield reduction in Danteswari and Samaleswari may be contributed by less reduction in PN, TDM and higher GF%.

### Correlation between grain yield and yield related traits

Under DS condition, DFF showed negative correlation with HI, PH, TN, GY, PN, FT% and GW but none of the correlation was significant except with HI (p<0.01). GY was positively associated with all the traits except DFF but showed a non-significant positive correlation with GW. TDM had a highly significant and positive correlation with GY (p<0.01), PN (p<0.01), FT% (p<0.01), TN (p<0.01) and PH (p<0.01). Similarly, GW showed non-significant correlation with all the traits except PN (p<0.05). Grain yield was significantly (P<0.01) correlated with DFF under WW condition and contributes 39.3% for grain yield, while it showed a non-significant positive correlation (p<0.2, r=0.237) under DS condition and contributes only 5.60% for grain yield. There was a positive significant correlation between grain yield and plant height (p<0.1) in WW condition while under DS condition, grain yield and plant height exhibited significant positive correlation (p <0.001). The tiller number, panicle number, total dry matter and grain filling percentage exhibited positively significant (p <0.001) correlation with grain yield both under WW and DS conditions (**Table 4**).

**Table – 4.**
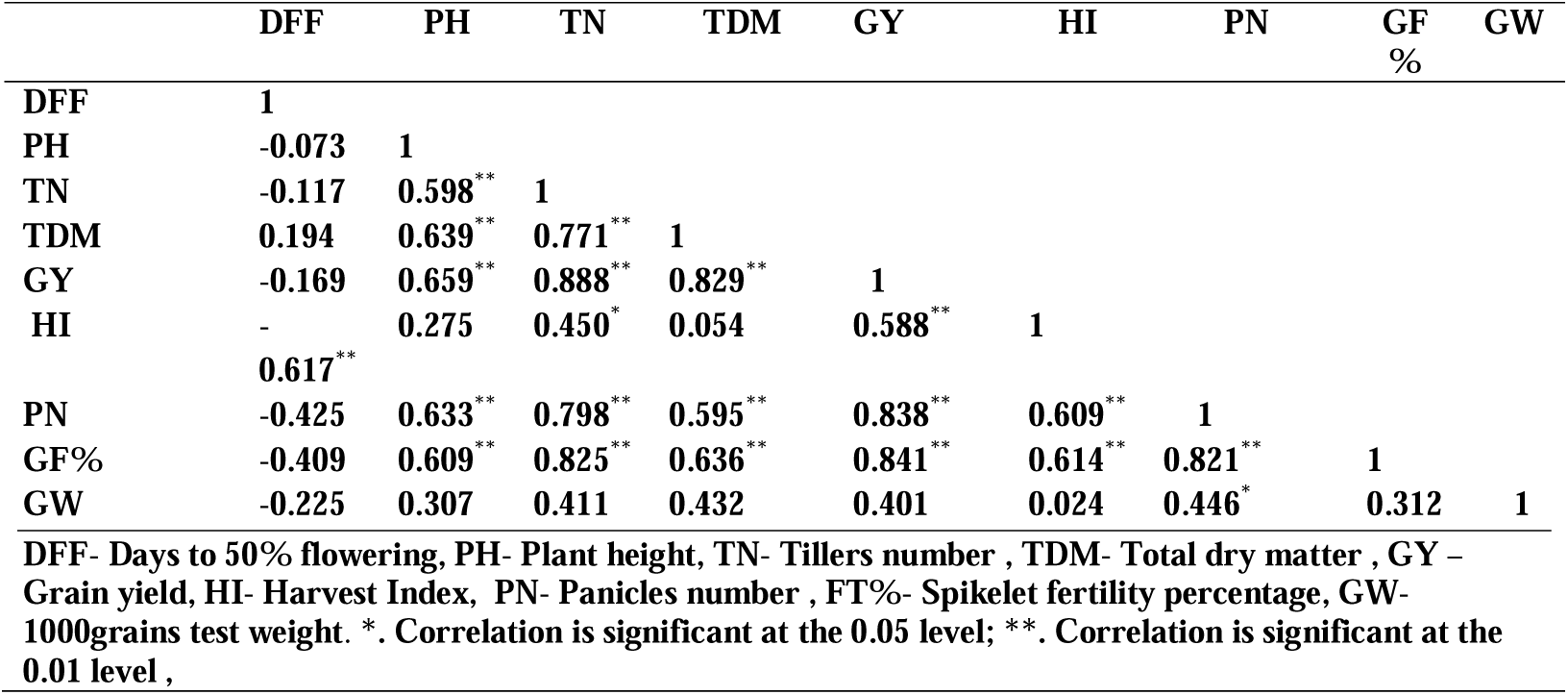
Correlation matrix of yield traits (pooled) under water stress condition

The Pearson correlation of the grain yield showed less positive (p>0.2) correlation with harvest index in WW condition and HI contributes 0.4%, while it was significantly positive (p<0.001) correlation between grain yield with harvest index under DS condition and HI contributes 66.40% to grain yield. There was a less positive (p>0.1) correlation of GW with grain yield and GW contributes only 1.61% to the grain yield under WW condition, while under DS condition, positive (p<0.05) correlation among GW and GY indicate GW contributes 17.58% to GY. Under WW condition TN, PN, TDM and GF% contributes 80.90%, 75.30%, 69.50% and 50.10% whereas under DS condition these parameters contributes 78.70%, 75.70%, 77.90% and 70.30% respectively towards grain yield (**Fig 8**).

**Fig 8.**
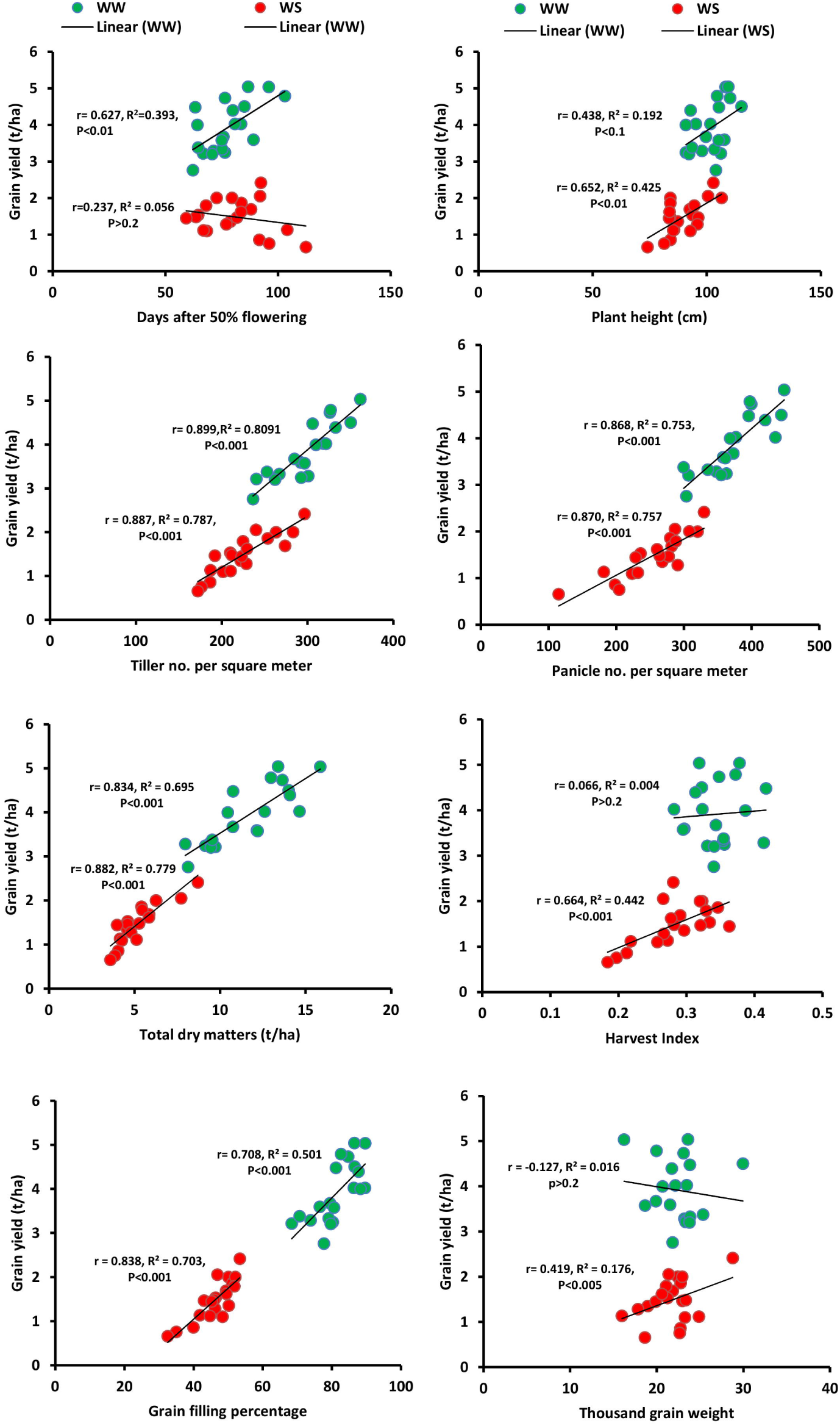
Correlation among the grain yield and yield related traits.

### Agglomerative Cluster analysis

Agglomerative Cluster analysis based on pearson correlation coefficient and unweight pair group analysis by considering grain yield and yield related traits, classified 20 genotypes (except Safri 17) into two main clusters I and II, at similarity coefficient of −0.635. The cluster I divided into 2 sub clusters: IA and IB, at similarity coefficient of 0.565 and the sub cluster IA and IB consist of 4 genotypes each. The cluster II divided into 2 sub clusters: IIA and IIB, at similarity coefficient of 0.365. The sub cluster IIA included 7 genotypes, and cluster IIB consist of 5 genotypes **(Fig 9)**. This pattern of clustering explained the existence of a significant amount of diversity among the genotypes.

**Fig 9.**
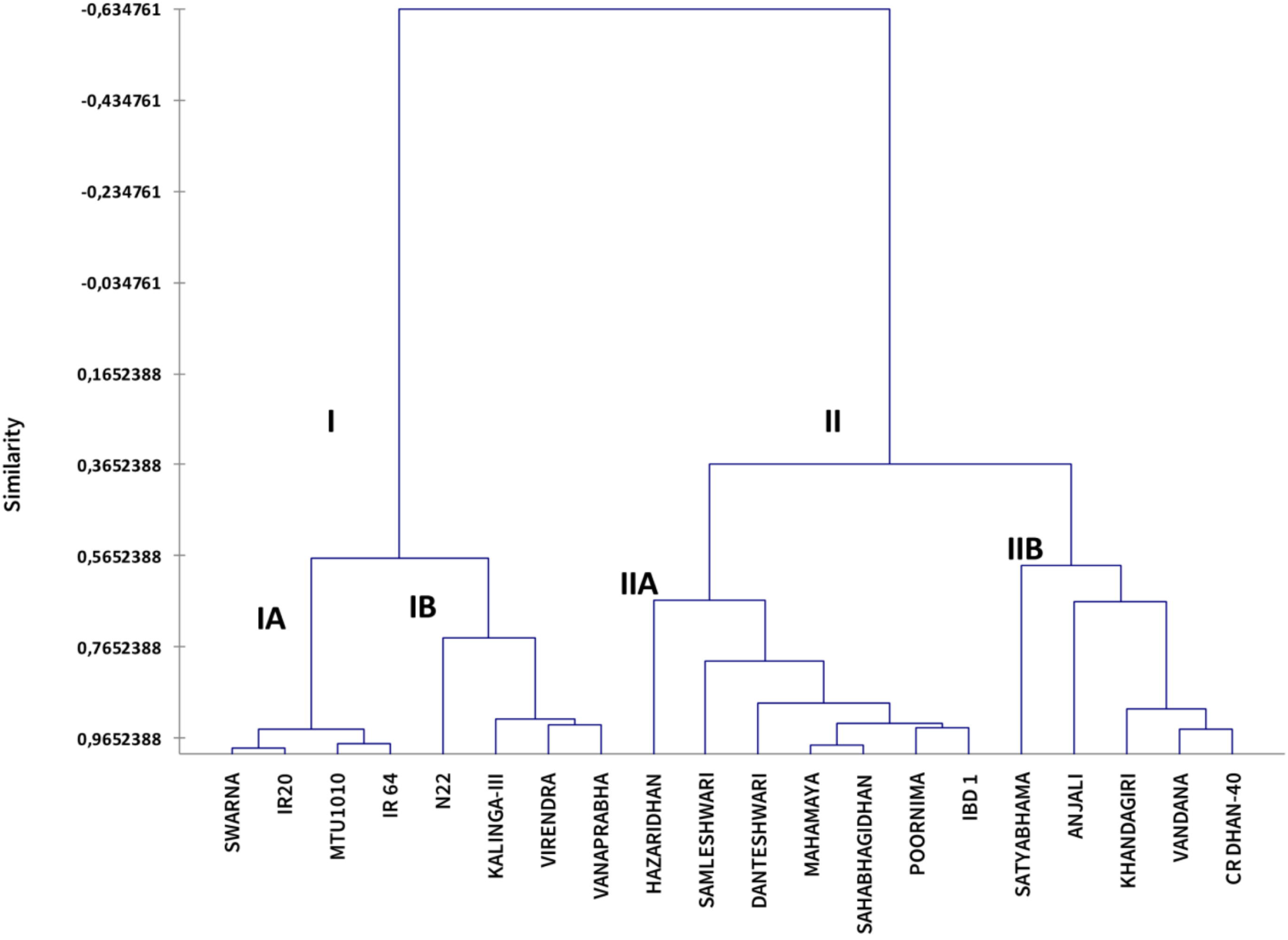
Dendrogram showing relationship among the 20 rice genotypes as revealed by Pearson correlation coefficient and Unweight pair group analysis based different yield traits.

Cluster IA included all the drought sensitive genotypes (Swarna, IR20, MTU 1010 and IR20) having longer DFF, and lowest tillers number, panicles number, total dry matters, grain yield, harvest index and grain filling percentage. Cluster IB consisted of genotypes (N22, Kalinga III, Virendra and Vanaprabha) having moderate to lower tiller number, panicles number, total dry matters and grain filling percentage. Clustered IIA is having genotypes (Hazaridhan, Samleshwari, Danteshwari, Mahamaya, Sahabhagidhan, Poornima and IBD-1) with high tiller number, panicle number, total dry matter, grain yield and grain filling percentage indicating drought tolerance and cluster IIB is characterized by the genotypes (Satyabhama, Anjali, Khandagiri, Vandana and CR Dhan 40) having moderate to lower value of DFF and higher to moderate value of plant height, panicles number and harvest index (**Fig 9)**.

## Discussion

In the present study attempt has been made to identify suitable genotype for rainfed upland condition of eastern India that mostly suffers from drought. Under drought stress condition, grain yield is determined by its phenological, physiological and yield traits (Barnaby *et al* 2019 In the present study stress was imposed both at vegetative and reproductive stage. Under drought, mean RWC decreased to 65.9% and LWP reached to −3.65 MPa. According to previous studies, LWP below −1.7 MPa affect plant growth (Santos *et al* 2018) which is supporting our results of decrease in plant height by 11.87% during stress period. Soil moisture tension during the stress was −47.83 kPa and −54.55 kPa in 2017 and 2018. Torres *et al*. (2018) reported similar results of occurring moderate to severe stress when soil moisture tension drops below −50 kPa. Drought stress at vegetative stage reduces water content and lower leaf water potential, leading to reduce turgor, stomatal conductance, and photosynthesis, and ultimately reduce grain yield (Akbarian *et al*., 2011; Amini *et al*., 2014). In our study, Mahamaya, Samaleswari, Poornima, IBD-1, Safri 17, Sahabhagidhan and N22 appears to be drought tolerant to the vegetative stage drought with SES drought score ‘1’at the end of the stress period and had early recovery after the stress was relieved. The identified genotypes also had high RWC (>70%) and LWP (>3.50 MPa) compared to susceptible genotypes like IR 20, IR 64 and MTU 1010 (Kumar *et al*., 2014). Drought score is used to measure the tolerance toward drought stress condition and reflects the extent of correlation of the plant tissue dehydration with its RWC (Ingram and Bartels, 1996; Cabuslay *et al*., 1999; 2002). The ability of the plant to recover after the drought relief is considered more crucial than drought tolerance (Fang and Xiong, 2015; Chen *et al*., 2016) which is important selection criterion for selecting drought tolerant genotypes. The seven identified genotypes had low drought score with higher recovery rate justifies their tolerance towards drought.

Though the genotypes have vegetative stage drought tolerance, they may not perform well in terms of grain yield production under drought (Swain *et al*., 2017). For effective screening, drought stress was imposed during flowering stage (Pantuwan *et al*., 2002) and to synchronize flowering staggered showing was adopted (Garrity and O’Toole, 1994). Out of seven genotypes showing tolerance at vegetative stage, only four genotypes (Mahamaya, Samleshswari, Sahabhagidhan and Danteswari) showed tolerance to reproductive stage drought stress. In this experiment, drought stress was severe enough to reduce grain yield to a greater extent. The result showed genotypic variation in grain yield between drought tolerant sensitive genotypes under a specific type of drought (Torres *et al*., 2018). Under drought stress, grain yield is one of the important selection criterions and it is determined by several phenological and yield attributes such as flowering duration, plant height, tiller numbers, panicle numbers, total dry matter, harvest index, grain filling percentage and 1000 grain weight. The most important trait contributing to drought tolerance in these four identified genotypes was higher leaf water potential that caused dehydration avoidance lead to higher biomass than IR 20 and IR 64 under stress. Due to high biomass, these four genotypes had better abilities to maintain a high growth rate under stress with less reduced PH, PN, and GF% under reproductive stage stress. According to Blum (2005), some genotypes have the mechanism of maintaining high plant water status despite of having more biomass and plant height due to less water using ability and more water absorbing capacity. This mechanism provided a good explanation for the significant positive correlation between TDM to GY and TDM to GF. From the above, the conclusion can be made that in eastern Asia, rainfed lowland rice is mostly a drought avoider, and produce higher grain yield under drought due to the ability of maintaining plant water status around flowering and grain filling (Fukai *et al*., 2009). Another mechanism that contributed to drought tolerance could be efficient partitioning partially resulting from shorter plant and less delay in DFF that resulted in higher GW and GF% under drought (Guan *et al*, 2010). Previous studies reported that increase in HI under drought is of critical importance for drought tolerance (grain yield) under terminal drought stress (Monneveux *et al*., 2008). But no such phenomenon was observed here, but a strong positive correlation (p<0.01) existed between HI with TDM and GY which indicates higher remobilization of assimilates to the grains from stems and leaves under drought (Kumar *et al*., 2006).

Drought escape or accelerated heading under drought, might contributed to drought tolerance. Our results showed that, N22 and Anjali flowered 5-7 days earlier compared to WW. Early flowering could partly responsible for the improved GF% and GW under DS because early flowering would allow plants to escape from severe terminal stress in rice (Xu *et al*., 2005). This drought escape by drought-induced accelerated heading in N22 and Anjali was expressed at the vegetative stage drought and had grain yield of >1.4 t/ha. It is important to mention here that the above discussed mechanisms function together and affected the same set of drought tolerance related traits but appeared to vary considerably depending on specific scenarios of drought stress. In our study significant variation in drought tolerance (in terms of grain yield) observed suggest that, for evaluation of genotypes should be performed under the type(s) of stress similar to the target environment. In the present study, VS had greater effects on traits like PH, PN and biomass (source supply), whereas RS more severely affected traits like GF% and GW that determine the sink size and partitioning. So, genotypes targeting for rainfed lowland where terminal drought is more frequent, should be evaluated under a severe RS. While selecting genotypes for rainfed upand, evaluation under both severe VS and RS would be required. Selection for grain yield under reproductive stress was practised due to the stress has much greater effect on the grain yield of rice (Serraj *et al*., 2009). However, visual selection for a large sink size under favourable conditions and higher source capacity (seedling vigour and tiller number) under VS and greater fertility under RS may help during large population screening (Blum, 2004).

## Conclusions

The present study identified some genotypes (Sahabhagidhan, Poornima, Vandana, and N22) that were tolerant to both vegetative as well as reproductive stage drought stress. Combining tolerance of both vegetative as well as reproductive-stage drought could be accomplished by performing separate trials for both the stresses that will help in the development of improved varieties with tolerance to multiple growth stages and help to maintain stable grain yield in rainfed ecosystem umder the prevailing unpredictable climatic situations.

## Declarations

### Ethics approval

NA

### Availability of data and material

All the required data are given in the manuscript

### Code availability

NA

## Acknowledgements

This study was financially supported by Jawaharlal Nehru Scholarship for doctoral studies, Jawaharlal Nehru memorial Fund, New Delhi.

